## Supplemental Figures S1-S6 and Tables S1-S5 for "The heat shock protein LarA activates the Lon protease at the onset of proteotoxic stress"

### Supplementary Figures and Tables

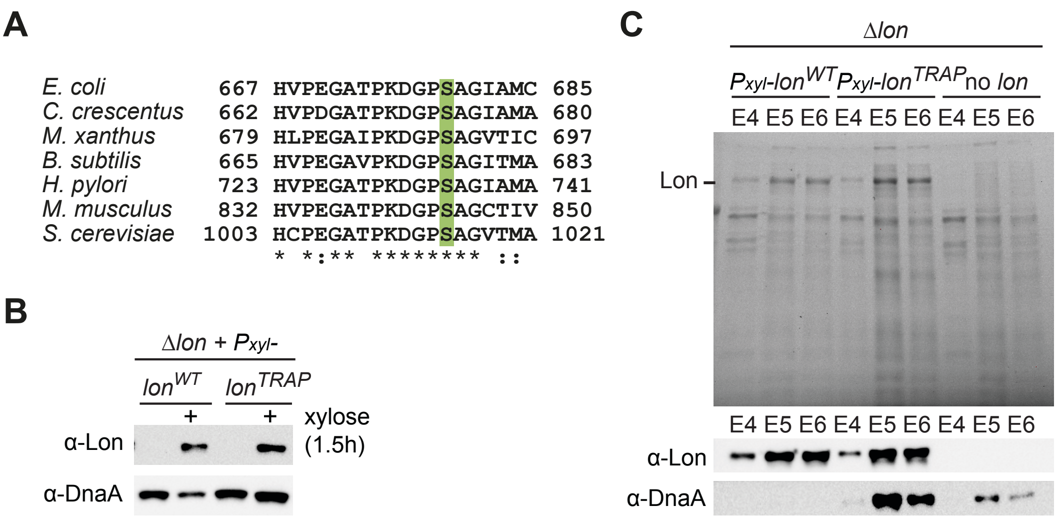

**Figure S1. A trapping approach allows co-purification of Lon-bound proteins.**

**(A)** Alignment of indicated amino acid residue sequences of Lon from different species (*Escherichia coli* [AJF45020.1], *Caulobacter crescentus* [WP_004615160.1], *Myxococcus xanthus* [WP_201424642.1], *Bacillus subtilis* [ASZ62291.1], *Helicobacter pylori* [GHQ92543.1], *Mus musculus* [NP_083058.2], *Saccharomyces cerevisiae* [GHM90547.1]) illustrating the conservation of the catalytic serine residue (highlighted in green) of the peptidase domain.

**(B)** Immunoblot analysis confirming the expression $Lon^{WT}$-Twin-Strep-tag and $Lon^{TRAP}$-Twin-Strep-tag in Δ*lon* cells for the protease trapping experiment after 1.5 hours of xylose-induction (upper panel). Levels of the known Lon substrate DnaA are negatively affected by expression of $Lon^{WT}$-Twin-Strep-tag but not of $Lon^{TRAP}$-Twin-Strep-tag, confirming its catalytic inactivity (lower panel).

**(C)** Stain-free SDS-PAGE gel analysis (upper panel, the bands corresponding to Lon derivatives are indicated at their height of migration) and immunoblot analysis (lower panel) of elution fractions 4-6 of the Twin-Strep-tag purification from strains described in Figure 1A, confirming the purification of Lon constructs and co-purification of DnaA specifically in Lon$^{TRAP}$-Twin-Strep-tag eluates. Representative of the two replicates.

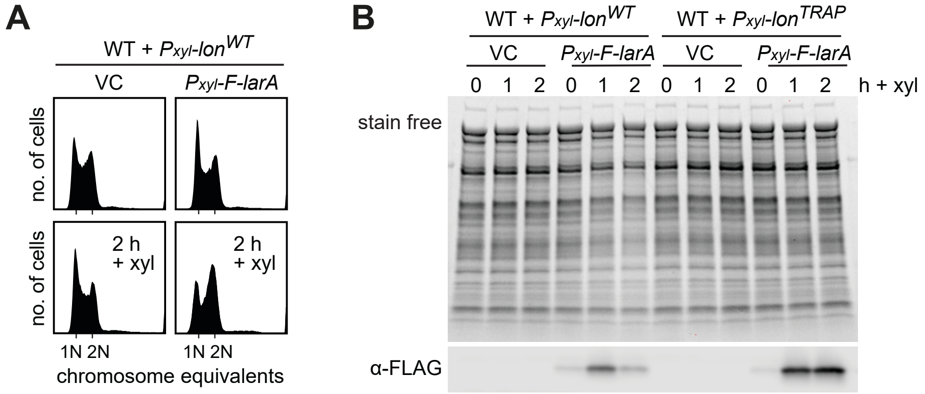

**Figure S2. Co-overexpression of *lon* and *larA* results in cell cycle arrest and reduced total protein content.**

**(A)** Flow cytometry analysis of wild type cells (WT) with $P_{xyl}$-*Lon*$^{WT}$-*Twin-Strep-tag* integrated on the chromosome and harboring an empty vector (VC) or a plasmid carrying $P_{xyl}$-*3xFLAG-larA* ($P_{xyl}$-*F-larA*). Samples were taken just before (0 h) and 2 hours after xylose addition to induce expression of *lon*$^{WT}$-*Twin-Strep-tag* and *F-larA* (2 h).

**(B)** Stain free SDS-PAGE gel (upper panel) and immunoblot analysis (lower panel) of extracts from wild type cells (WT) with either chromosomally integrated $P_{xyl}$-*Lon*$^{WT}$-*Twin-Strep-tag* or $P_{xyl}$-*Lon*$^{TRAP}$-*Twin-Strep-tag* and harboring an empty vector (VC) or a plasmid carrying $P_{xyl}$-*3xFLAG-larA* ($P_{xyl}$-*F-larA*). Samples were taken at time point 0 as well as 1 and 2 hours after xylose addition to induce expression of *lon*$^{WT}$-*Twin-Strep-tag* or *lon*$^{TRAP}$-*Twin-Strep-tag* as well as of *F-larA* where indicated.

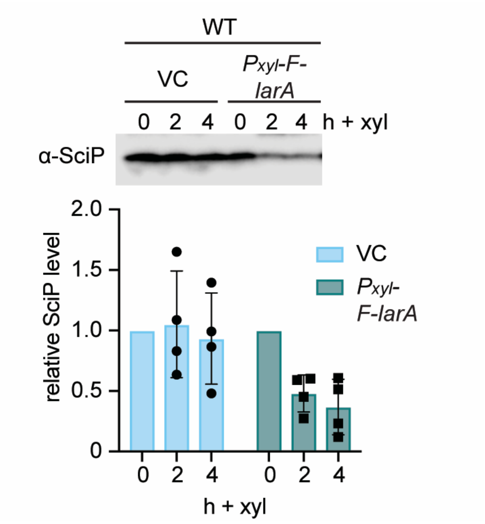

**Figure S3. LarA overexpression results in reduced SciP levels.**

Immunoblot analysis of SciP levels in wild type cells (WT) harboring an empty vector (VC) or a plasmid carrying $P_{xyl}$-*3xFLAG-larA* ($P_{xyl}$-*F-larA*), respectively. Samples were taken at the indicated time points after xylose addition to induce expression of *F-larA* where indicated. Graph shows the relative SciP levels from 4 biological replicates as well as the means, error bars represent standard deviations.

**A**

```
Methylobacterium      ----MTVLNPSPLTPADLDPAEFQAADLAALGEGHIAYVRPIRSDEVRRFFPQA-
Bradyrhizobium        --------MSEGHVAFEYEAKNVSPETLATLGEGHIAYVKQIRSEDVPDLFPEA-
Agrobacterium         -------------MLLKEAHARLTQSQLAHIGEGEVGYIRKIRAEEVSRCFPEA-
Shinella              -------------MGLKSVTPTVSKADLAHLGAGEVGYIRKMKADEVSRVFPEA-
Bartonella            ---------------MGEHKQNLSLQDMTHSDTGQIAYLKKVCTDDLAKNFPDL-
Mesorhizobium         MALSGVLAPRRLEMTRTDENITMTSGEFAHLGEGSVAYLRKVSSDELLGRFPNI-
Brucella              ------------MEAKTMNKHVFTDNEFAHLGEGEIAYVRKIRSDDLARSFPSL-
Rhodobacterales       ---------------------MHTPFDFKDAGSRTV-YVKAVAVADLPKEVQAS-
Rhodobacterales       ---------------------METPYEFEEDGPDRIVYIRSVEVAELPEDLRDQ-
Rhodobacter           ---------------------MNTPFDFPE-GAGRLVYVRPVPVAELPEDVQSM-
Roseobacter           ---------------------MQTRYDFGADEGRPIVYVRPVHVADLPEDMREQ-
Oceanicaulis          ---------------------MDTETELMLDTTPTV-YVRTIDAAEIMEQAANA-
Caulobacter           ------------MTPSTQHSRPLTSEAFAALGAPALVYVRPIKAAEILADAPEG-
Phenylobacterium      -------------MEVVMMTPVLSTEAFAALGAPHLVYVRPVKAAEILASVPSSQ
Micavibrio            -----MTDCKTTDCKSASALRDISAQDFLGFGVQHVAYVRPA-------------
Magnetospirillum      ------------MQDLSTLLYPMSASDFASWGMPELAFVKRV-------------
Elioraea              --------MNTQKTRGQVSIPVPTRQQLAELGLGGLAYVRAVVVD----------
Kiloniella            ----------------------MTEQDMTALGLEAVAYVKAVDVD----------
                                                 :       : ::.

Methylobacterium      LTPGLDLFALLSASGAPIMLADSRDVVLANAAAHDLQPVSLH
Bradyrhizobium        IAPGLKLFALHAADGTPIMLTDSREAAVANAWSNELQAVSVH
Agrobacterium         LDPQLDLWALFGADGTPILLTDNRSSTFFKAAEDDLKTVSLH
Shinella              IDPSLDLWALFGADGTPILLTDNRSSTFYKAAEDDLRTVSLH
Bartonella            MTPGITVWALFGETGYPIILSDERSIALAGANEHELQIVTLH
Mesorhizobium         IAPGLELWALFAANGQPILLSDARDRALAGAMENDLTTVAIH
Brucella              IAPGLELWALFGASGEPIVLSDVRATALQGAHDHELRTVTLH
Rhodobacterales       AGGREQLYAVHDAEGGQLALVADRSLAFVLARQNDMTPVPVH
Rhodobacterales       AEGRDQIYAVHSEDGERLALVRDRAMAFALARSNDLAPVAVH
Rhodobacter           ARGLDHIYALHGADGERLALVRDRHLAFVLARQNDLAPVNVH
Roseobacter           AMGSTTLYALHSAEGERLALVRDRRLAFVLARQNNMNPVAVH
Oceanicaulis          LDSDATLYAIHDENGQRMAVFSDREAAFAAARDHGAEPVSVH
Caulobacter           LSPDQTLYAVCRADGERLAVLIDRDTAIAAALAHELAPVSVH
Phenylobacterium      LDPDQTLYAVHRADGERLAVLTDRESAVAAALAHELAPVSVH
Micavibrio            ILNGQKVFSIHAADGTPLGYHDNFDAALVLTRQNDLEPVRVQ
Magnetospirillum      VENSQVSWTIHSADGTAIGVAPSRDLAFAAIIQHELEPVSVH
Elioraea              ---GQSAFAIFGADGQQIGLAPDRALADAAIVQHEMVPVSVH
Kiloniella            ---GEACFGIFAADGTEITVVTNRDVAFAAVRQQDLEPLSVH
                             : :    *  :        .           : ::
```

**B**

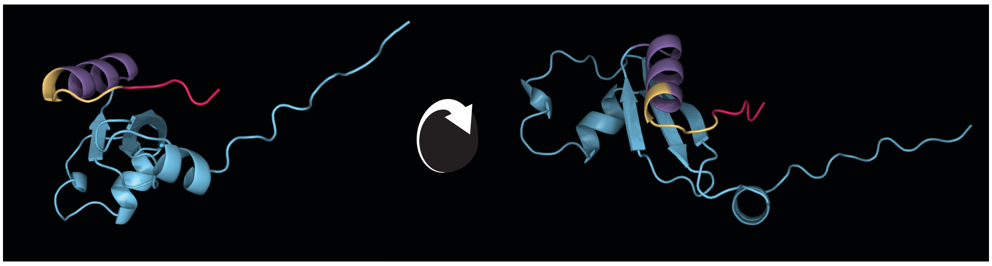

**Figure S4. The C-terminus of LarA is highly conserved and folds into a α-alpha-helix followed by a short unstructured region.**

**(A)** Alignment of LarA amino acid residue sequences from different species illustrating the high level of conservation of the C-terminal 5 amino acids. *Caulobacter vibrioides* [YP_002519080.1], *Rhodobacter sphaeroides* [WP_002722317.1], *Brucella metlitensis* [WP_002968114.1], *Agrobacterium tumefaciens* [WP_006310223.1],

*Bartonella Henselae* [WP_011180040.1], *Kiloniella laminariae* [WP_157230811.1], *Micavibrio aeruginosavorus* [WP_014101741.1], *Elioraea tepidiphila* [WP_019014739.1], *Magnetospirillum gryphiswaldense* [WP_024081410.1], Rhodobacterales bacterium [WP_008556561.1], *Methylobacterium radiotolerans* [WP_012320107.1], *Bradyrhizobium japonicum* [WP_018645064.1], *Shinella* sp. DD12 [WP_024270222.1], *Phenylobacterium zucineum* [WP_012520744.1], *Oceanicaulis* sp. HTCC2633 [WP_009802573.1], Rhodobacterales bacterium HTCC2150 [GenBank: EBA04284.1], *Roseobacter* sp. AzwK-3b [ZP_01904599.1], *Mesorhizobium* [WP_010911485.1]. Alignment was created using EMBL-EBI Multiple Sequence alignment MUSCLE [1].

**(B)** LarA structure predictions by Colabfold [2]. The C-terminal amino acid residues are predicted to form an α-helix (residues 71 to 81) followed by an unstructured region (residues 82 to 89). Together, those correspond to the C-terminal 20 amino acids analysed in this study. The amino acid residues that were removed in the LarA$^{\Delta 20}$, LarA$^{\Delta 10}$, LarA$^{\Delta 5}$ mutants are colored in magenta, yellow and red (Δ20), yellow and red (Δ10) and red only (Δ5), respectively.

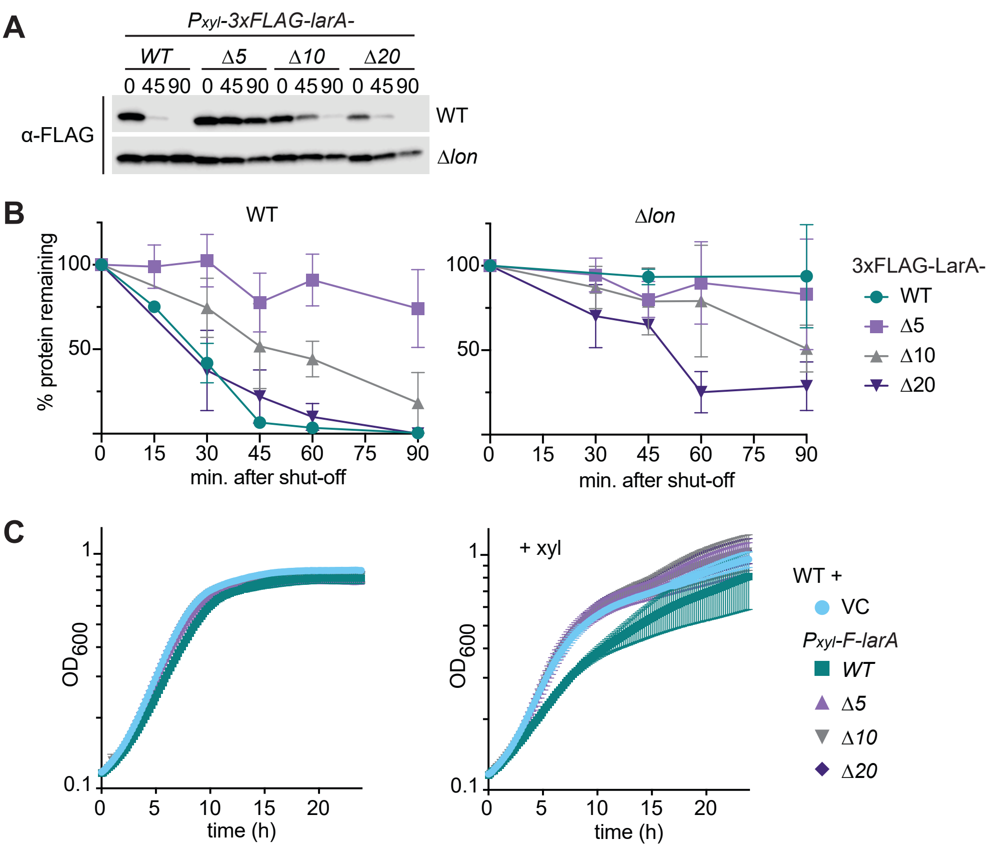

**Figure S5. The C-terminal amino acids of LarA are required for *in vivo* LarA degradation and LarA-dependent effects on growth.**

**(A)** Immunoblot analysis of stability of 3xFLAG-LarA (F-LarA; WT) and F-LarA variants in wild type (upper panel) and Δ*lon* (lower panel) cells harboring plasmids carrying either *$P_{xyl}$-3xFLAG-larA* (*WT*), *$P_{xyl}$-3xFLAG-larA$^{\Delta 5}$* (Δ*5*), *$P_{xyl}$-3xFLAG-larA$^{\Delta 10}$* (Δ*10*), or *$P_{xyl}$-3xFLAG-larA$^{\Delta 20}$* (Δ*20*), respectively. Expression of F-LarA variants was induced by addition of xylose for 1 hour prior to addition of chloramphenicol to shut-off protein synthesis and samples to assess protein stability were subsequently taken at the indicated time points. **(B)** Graphs show the means of F-LarA variant levels in wild type (left hand side) and Δ*lon* cells (right hand side) from two biological replicates, error bars represent standard deviations.

**(C)** Growth experiment using a plate reader to assess $OD_{600}$ over 24 hours. Wild type cells (WT) harboring an empty vector (VC) or plasmids harboring either *$P_{xyl}$-3xFLAG-*

*larA* (*WT*), *$P_{xyl}$-3xFLAG-larA*$^{\Delta 5}$ (Δ*5*), *$P_{xyl}$-3xFLAG-larA*$^{\Delta 10}$ (Δ*10*), or *$P_{xyl}$-3xFLAG-larA*$^{\Delta 20}$ (Δ*20*), respectively, were grown under non-inducing conditions (left panel) or with xylose, i.e., $P_{xyl}$-inducing conditions (+ xyl; right panel). All growth curves display the means of three biological replicates; error bars represent standard deviations.

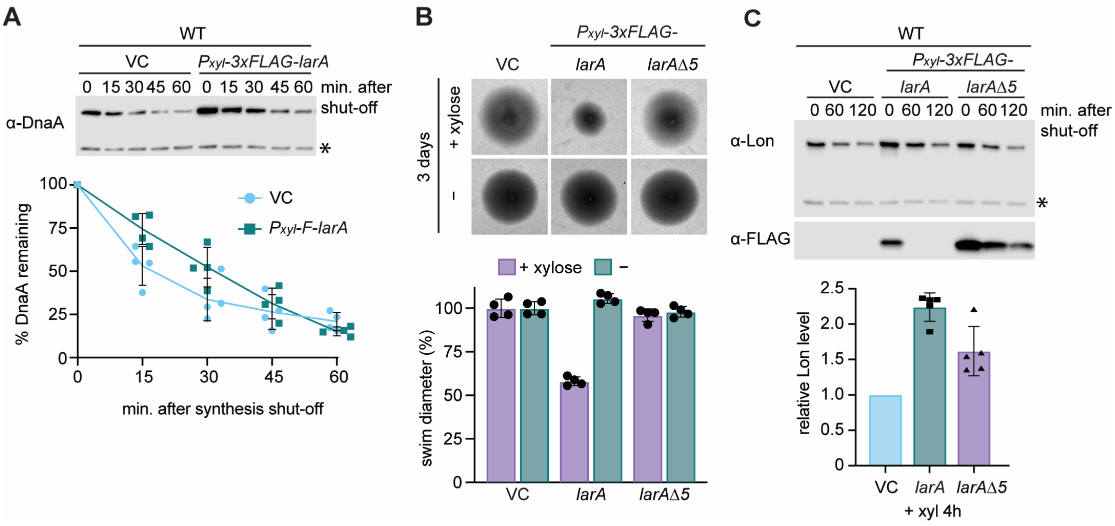

**Figure S6. Effects of *larA* overexpression on DnaA abundance and stability, soft-agar motility as well as Lon abundance.**

**(A)** Immunoblot analysis of DnaA levels in wild type cells (WT) harboring an empty vector (VC) or a plasmid carrying $P_{xyl}$-*3xFLAG-larA* ($P_{xyl}$-*F-larA*). Expression of F-LarA was induced by addition of xylose for 1 hour prior to addition of chloramphenicol to shut-off protein synthesis and samples to assess DnaA stability were subsequently taken at the indicated time points. Graph shows the means of DnaA levels from four biological replicates, error bars represent standard deviations.

**(B)** Motility assay in PYE soft agar containing gentamycin of wild type cells harboring the empty vector (VC) or overproducing 3xFLAG-tagged LarA ($P_{xyl}$-*3xFLAG-larA*) or F-LarA$^{\Delta5}$ ($P_{xyl}$-*3xFLAG-larA*$^{\Delta5}$), respectively, by xylose induction (+ xylose) in comparison to non-inducing conditions (−). The graph shows the relative swim diameters from four biological replicates, means (means of VC were set to 100%) and standard deviations are indicated.

**(C)** Immunoblot analysis of Lon levels (upper panel) in wild type cells (WT) harboring an empty vector (VC), a plasmid carrying $P_{xyl}$-*3xFLAG-larA* or $P_{xyl}$-*3xFLAG-larA*$^{\Delta5}$. Expression of *F-larA* and *F-larA*$^{\Delta5}$ was induced by addition of xylose for 4 hours prior to addition of chloramphenicol to shut-off protein synthesis and samples to assess Lon stability were subsequently taken at the indicated time points. Graph shows the relative Lon steady state levels after 4 hours of xylose induction from five biological replicates, means are indicated and error bars represent standard deviations.

**Table S1. Kinetic parameters of LarA-activated His-SciP degradation, related to Figure 3C.**

| | $V_b$ ($min^{-1}$ $Lon_6^{-1}$) | $V_{max}$ ($min^{-1}$ $Lon_6^{-1}$) | $V_i$ ($min^{-1}$ $Lon_6^{-1}$) | $n$ | $K_a$ (µM) | $K_i$ (µM) |
|---|---|---|---|---|---|---|
| Value | 1.5 ± 0.3 | 9.0 ± 3.0 | 6.8 ± 0.5 | 1.5 ± 0.5 | 0.3 ± 0.2 | 3.6 ± 5.9 |
| p-value | 2.01e-05 *** | 0.00625 ** | 1.41e-11 *** | 0.00901 ** | 0.04548 * | 0.55446 |

Parameters show value ± standard error determined by fitting a formula derived from (7) of [3] to at least 3 experimentally determined degradation rates of His-SciP at various LarA concentrations. $V_b$: basal degradation without LarA; $V_{max}$: theoretical maximum degradation rate; $V_i$: max degradation at full inhibition; $n$: Hill constant; $K_a$: concentration with half-maximum activation; $K_i$: concentration with half-maximum inhibition. Asterisks indicate significance levels: *** $p < 0.001$, ** $p < 0.01$, * $p < 0.05$

**Table S2. Kinetic parameters of Lon-dependent His-SciP degradations in the presence or absence of LarA, related to Figure 3D.**

| **Substrates** | $V_{max}$ ($min^{-1}$ $Lon_6^{-1}$) | $K_m$ (µM) | **Hill constant** $n$ | $k_{cat}$ ($min^{-1}$ $Lon_6^{-1}$) | **catalytic efficiency** ($µM^{-1}$ $min^{-1}$ $Lon_6^{-1}$) |
|---|---|---|---|---|---|
| His-SciP | 7.6 ± 1.7 *** | 9.3 ± 4.0 * | – | 151.1 | 16.2 |
| His-SciP + LarA | 15.0 ± 0.8 *** | 1.8 ± 0.2 *** | 1.9 ± 0.4 *** | 300.5 | 166.1 |

Values for $V_{max}$, $K_m$ and $n$ represent parameters ± standard error determined by fitting the Michaelis-Menten and Hill equations to experimentally determined degradation rates with at least 3 independent measurements per concentration. $k_{cat}$ and catalytic efficiency were calculated based on those values. Asterisks indicate significance levels: *** $p < 0.001$, ** $p < 0.01$, * $p < 0.05$

**Table S3. Strains used in this study.**

| ***Caulobacter crescentus* strains** | | | |
|---|---|---|---|
| **Name** | **Genotype** | **Marker** | **Reference** |
| NA1000 (CB15N) | WT, synchronizable derivative of wild-type CB15 | | Provided by Michael Laub |
| LS2382 | Δ*lon (*NA1000 *lon*::*Ω)* | spec$^R$ | [4] |
| KJ546 (=KG329) | Δ*lon* (*lon*::*Ω* re-introduced into NA1000 by phage transduction) | spec$^R$ | |
| KJ1066 (=DJO002) | Δ*lon* $P_{xyl}$-*lon*$^{WT}$-*TwinStrep-tag* | kan$^R$ | This study |
| KJ1067 (=DJO003) | Δ*lon* $P_{xyl}$-*lon*$^{TRAP}$-*TwinStrep-tag* | kan$^R$ | This study |
| KJ1068 (=DJO004) | WT $P_{xyl}$-*lon*$^{WT}$-*TwinStrep-tag* | kan$^R$ | This study |
| KJ1069 (=DJO005) | WT $P_{xyl}$-*lon*$^{TRAP}$-*TwinStrep-tag* | kan$^R$ | This study |
| KJ1070 (=DJO006) | Δ*larA(CCNA_03707)* | tet$^R$ | This study |
| ***Escherichia coli* strains** | | | |
| **Name** | **Genotype** | **Marker** | **Reference** |
| DH5α | General cloning strain | | Invitrogen |
| BL21-SI/ pCodonPlus | Salt-inducible BL21(DE3) strain for protein expression | chlor$^R$ | Provided by Claes Andréasson |
| BL21(DE3)/ pLysS | Protein expression strain | chlor$^R$ | Lab collection |
| ER2566 | Lon-deficient B strain for protein expression | | Provided by Peter Chien (originally from NEB) |

#### Table S4. Plasmids used in this study.

| Name | Description | Marker | Reference |
|---|---|---|---|
| pBX-MCS-2 | High copy, xylose-inducible expression | kan$^{R}$ | [5] |
| pBX-MCS-4 | High copy, xylose-inducible expression | gent$^{R}$ | [5] |
| pDJO26 | pBX-MCS-2 containing *lon*$^{WT}$-*TwinStrep-tag* | kan$^{R}$ | This study |
| pDJO40 | pBX-MCS-2 containing *lon*$^{TRAP}$-*TwinStrep-tag* | kan$^{R}$ | This study |
| pDJO145 | pBX-3xFLAG-4 | gent$^{R}$ | [6] |
| pDJO305 | pBX-MCS-4 containing $P_{xyl}$-*larA-3xFLAG* | gent$^{R}$ | This study |
| pDJO307 | pBX-MCS-4 containing $P_{xyl}$-*3xFLAG-larA* | gent$^{R}$ | This study |
| pDJO374 | pBX-MCS-4 containing $P_{xyl}$-*3xFLAG-larAΔ5* | gent$^{R}$ | This study |
| pDJO377 | pBX-MCS-4 containing $P_{xyl}$-*3xFLAG-larAΔ10* | gent$^{R}$ | This study |
| pDJO380 | pBX-MCS-4 containing $P_{xyl}$-*3xFLAG-larAΔ20* | gent$^{R}$ | This study |
| pDJO451 | pBX-MCS-4 containing $P_{xyl}$-*3xFLAG-larA-H89D* | gent$^{R}$ | This study |
| pDJO460 | pBX-MCS-4 containing $P_{xyl}$-*3xFLAG-larA-V88D-H89D* | gent$^{R}$ | This study |
| pDJO461 | pBX-MCS-4 containing $P_{xyl}$-*3xFLAG-larA-V86A-V88A* | gent$^{R}$ | This study |
| pML1716-*lon* (KJ600) | pML1716 containing *lon* | chlor$^{R}$ | [7] |
| pXCHYN-2 | Plasmid to integrate at *xylX* locus | kan$^{R}$ | [5] |
| pDJO67 | pXCHYN-2 containing *lon*$^{WT}$-*TwinStrep-tag* | kan$^{R}$ | This study |
| pDJO70 | pXCHYN-2 containing *lon*$^{TRAP}$-*TwinStrep-tag* | kan$^{R}$ | This study |
| pNPTS138 | Integrating vector for two-step recombination | kan$^{R}$ | Lab collection, provided by Michael Laub |
| pDJO404 | pNPTS138-*UHR-tet-DHR(larA)*, generation of a *tet*$^{R}$-marked deletion of *larA*(*CCNA_03707)* | tet$^{R}$, kan$^{R}$ | This study |
| pBAD33-Lon6his | pBAD33 derived vector for L-arabidose induced Lon-6xHis expression | chlor$^{R}$ | Provided by Peter Chien |
| pSUMO-YHRC | Plasmid for protein expression using $P_{T7}$ with an N-terminal 6xHis-SUMO tag; RRID:Addgene_54336 | kan$^{R}$ | [8] |
| pHis-SciP | pET-6xHis-sciP | amp$^{R}$ | [9] |
| pMF65-c5 | pSUMO-YHRC containing *6xHis-SUMO-larA* | kan$^{R}$ | This study |
| pMF89-c2 | pSUMO-YHRC containing *6xHis-SUMO-larAΔ5* (Δ85-89) | kan$^{R}$ | This study |
| pMF82 | pSUMO-YHRC containing *6xHis-SUMO-larA-V86A-V88A* | kan$^{R}$ | This study |
| pMF81-c2 | pSUMO-YHRC containing *6xHis-SUMO-larA-H89D* | kan$^{R}$ | This study |

| **Name** | **Description** | **Marker** | **Reference** |
|---|---|---|---|
| pMF88-c4 | pSUMO-YHRC containing *6xHis-SUMO-larA-V88D-H89D* | $kan^R$ | This study |
| pMF58-c4 | pSUMO-YHRC containing *6xHis-SUMO-fliX* | $kan^R$ | This study |
| pSH21-6xHis-titinI27-β20 | pSH21 containing coding sequence for N-terminal 6xHis tagged human titinI27 domain with β-galactosidase degron at the C-terminus (*6xHis-titinI27-β20*) | $amp^R$ | [10] |
| pAK002 | pSH21 containing *6xHis-titinI27-larA5* | $amp^R$ | This study |
| pAK003 | pSH21 containing *6xHis-titinI27-larA10* | $amp^R$ | This study |
| pAK004 | pSH21 containing *6xHis-titinI27-larA20* | $amp^R$ | This study |
| pAK005 | pSH21 containing *6xHis-titinI27-larA5-V88D-H89D* | $amp^R$ | This study |
| pAK006 | pSH21 containing *6xHis-titinI27-LarA5-V86A-V88A* | $amp^R$ | This study |

**Table S5. Oligonucleotides used in this study.**

| Name | Sequence (5'-3') | Reference |
|---|---|---|
| OAK057 | CGCGGATCCCTAGTGGACAGACACCGGACTAGTCC | This study |
| OAK058 | GGACTAGTCCGGTGTCTGTCCACTAGGGATCCGCG | This study |
| OAK059 | CGCGGATCCCTAGTGGACAGACACCGGCGCCAGCTCATGGGCACTAGTCC | This study |
| OAK060 | GGACTAGTGCCCATGAGCTGGCGCCGGTGTCTGTCCACTAGGGATCCGCG | This study |
| OAK061 | CGCGGATCCCTAGTGGACAGACACCGGCGCCAGCTCATGGGCGAGGGCGGCGGCGATCGCGGTGTCGCGGTCACTAGTCC | This study |
| OAK062 | GGACTAGTGACCGCGACACCGCGATCGCCGCCGCCCTCGCCCATGAGCTGGCGCCGGTGTCTGTCCACTAGGGATCCGCG | This study |
| OAK079 | CGCGGATCCCTAGTCGTCAGACACCGGACTAGTCC | This study |
| OAK080 | GGACTAGTCCGGTGTCTGACGACTAGGGATCCGCG | This study |
| OAK081 | CGCGGATCCCTAGTGGGCAGACGCCGGACTAGTCC | This study |
| OAK082 | GGACTAGTCCGGCGTCTGCCCACTAGGGATCCGCG | This study |
| oDJO13 | ATGGTCGTCTCCCCAAAACTC | This study |
| oDJO15 | AGCCCGGGGGATCCACTAGTTC | This study |
| oDJO16 | GAGTTTTGGGGAGACGACCATATGTCCGAACTACGTACGCTTCCTG | This study |
| oDJO171 | GCTCGAGTTTTGGGGAGACGACCATATGACGCCCAGCACCCAACAC | This study |
| oDJO172 | CACCGTCATGGTCTTTGTAGTCCATATGGTGGACAGACACCGGCGCCAG | This study |
| oDJO173 | GACTACAAGGACGACGACGACAAGGGTACCATGACGCCCAGCACCCAACAC | This study |
| oDJO174 | AGTGGATCCCCCGGGCTGCAGTTAGGTACCCTAGTGGACAGACACCG | This study |
| oDJO18 | GAACTGCGGGTGGCTCCAGCTAGCGTGCGTCAGCATGGCGTCGCTG | This study |
| oDJO182 | AGTGGATCCCCCGGGCTGCAGTTAGGTACCCTACGCCAGCTCATGGGCGAG | This study |
| oDJO183 | AGTGGATCCCCCGGGCTGCAGTTAGGTACCCTAGAGGGCGGCGGCGATC | This study |
| oDJO184 | AGTGGATCCCCCGGGCTGCAGTTAGGTACCCTAGATCAGGACCGCGAG | This study |
| oDJO185 | CAATTGAAGCCGGCTGGCGCCAAGCTTCGGTCTTCACGAACGAAGTCGC | This study |
| oDJO186 | GTATAGGAACTTCATGAATTCGATATCAAGCTTATCGATACCGGGTGCTGGGCGTCATAGGACC | This study |
| oDJO187 | GTTCCTATACTTTCTAGAGAATAGGAACTTCTTGAATTCCTGCAGGAGCTGGCGCCGGTGTCTGTCC | This study |
| oDJO188 | CCTGTACATCCGGAGACGCGTCACGGCCGAAGCTAGCGAATTCCAGTCGCTGGAGCGCCAAGG | This study |
| oDJO193 | CTCCTCTTGAACCGAC | This study |
| oDJO194 | GAACAGCGTGTTTCG | This study |
| oDJO197 | AGTGGATCCCCCGGGCTGCAGTTAGGTACCCTAGTCGACAGACACCG | This study |

| Name | Sequence (5'-3') | Reference |
|---|---|---|
| oDJO198 | AGTGGATCCCCCGGGCTGCAGTTAGGTACCCTAGTCGTCAGACACCG | This study |
| oDJO199 | AGTGGATCCCCCGGGCTGCAGTTAGGTACCCTAGTGGGCAGACGCCGG | This study |
| oDJO20 | GAACTAGTGGATCCCCCGGGCTTTAGGCGCCTTTTTCGAACTGC | This study |
| oDJO21 | TGGCTCCACGATCCACCTCCCGATCCACCTCCGGAACCTCCACCTTTCTCGAACTGCGGGTGGCTCCAGC | This study |
| oDJO22 | CTAGAACTAGTGGATCCCCCGGGCTTTAGGCGCCTTTTTCGAACTGCGGGTGGCTCCACGATCCACCTCC | This study |
| oDJO23 | CACGCCCAAGGATGGTCCGGCTGCAGGCATCGCCATGGCCTTGG | This study |
| oDJO24 | CCAAGGCCATGGCGATGCCTGCAGCCGGACCATCCTTGGGCGTG | This study |
| oDJO25 | GCGTAACGTTCGAATTCTCCGGAGCTCTTAGGCGCCTTTTTCGAACTGC | This study |
| OFS25 | CGGTATCGATAAGCTTGATATCGAATTCATGAAGTTCCTATAC | Lab collection |
| OFS26 | CTGCAGGAATTCAAGAAGTTCCTATTCTCTAGAAAGTATAGGAAC | Lab collection |
| OFS932 | CTCGAGTTTTGGGGAGACGACCATATGTCCGAACTACGTACGCTTCCTGTC | Lab collection |
| oMJF106 | GTGCGGCCGCAAGCTTGTCGACGGAGCTCGAATTCGGATCCTAATCGTCAGACACCGGCGCCAGCTC | This study |
| oMJF107 | GTGCGGCCGCAAGCTTGTCGACGGAGCTCGAATTCGGATCCTACGCCAGCTCATGGGCGAG | This study |
| oMJF34 | CCCACCAATCTGTTCTCTGTG | [6] |
| oMJF36 | CATGCATCATCAGGAGTACGG | [6] |
| oMJF37 | GATCCGAATTCGAGCTCC | [6] |
| oMJF38 | GAATTTATGCCTCTTCCGACC | [6] |
| oMJF47 | AGAGAACAGATTGGTGGGATGAAGGTTTCCAGCACG | This study |
| oMJF48 | GGAGCTCGAATTCGGATCTGCTATCCGGCCCTG | This study |
| oMJF67 | TAACGATATTATTGAGGCTCACAGAGAACAGATTGGTGGGATGACGCCCAGCACCCAACAC | This study |
| oMJF68 | GTGCGGCCGCAAGCTTGTCGACGGAGCTCGAATTCGGATCCTAGTGGACAGACACCG | This study |
| oMJF96 | GTGCGGCCGCAAGCTTGTCGACGGAGCTCGAATTCGGATCCTAATCGACAGACACCGGCGCCAGC | This study |
| oMJF97 | GTGCGGCCGCAAGCTTGTCGACGGAGCTCGAATTCGGATCCTAGTGTGCAGATGCCGGCGCCAGCTCATGGGCGAG | This study |
| RecUni-1 | ATGCCGTTTGTGATGGCTTCCATGTCG | [5] |
| RecXyl-2 | TCTTCCGGCAGGAATTCACTCACGCC | [5] |
| T7 | TAATACGACTCACTATAGGG | common primer |
| T7 terminator | GCTAGTTATTGCTCAGCGG | common primer |

### Supplementary Datasets

**Dataset S1.** Proteomics-based identification of Lon-bound proteins, related to Figure 1.

**Dataset S2.** Quantitative proteomics analysis of LarA overexpressing cells, related to Figure 6.

### References

1 Madeira, F. *et al.* The EMBL-EBI search and sequence analysis tools APIs in 2019. *Nucleic Acids Res* **47**, W636-W641, doi:10.1093/nar/gkz268 (2019).

2 Mirdita, M., Ovchinnikov, S., Steinegger, M. ColabFold: Making protein folding accessible to all. *bioRxiv* **8** (2021).

3 Walsh, R., Martin, E. & Darvesh, S. A versatile equation to describe reversible enzyme inhibition and activation kinetics: modeling beta-galactosidase and butyrylcholinesterase. *Biochim Biophys Acta* **1770**, 733-746, doi:10.1016/j.bbagen.2007.01.001 (2007).

4 Wright, R., Stephens, C., Zweiger, G., Shapiro, L. & Alley, M. R. Caulobacter Lon protease has a critical role in cell-cycle control of DNA methylation. *Genes & development* **10**, 1532-1542 (1996).

5 Thanbichler, M., Iniesta, A. A. & Shapiro, L. A comprehensive set of plasmids for vanillate- and xylose-inducible gene expression in Caulobacter crescentus. *Nucleic Acids Res* **35**, e137, doi:10.1093/nar/gkm818 (2007).

6 Omnus, D. J., Fink, M. J., Szwedo, K. & Jonas, K. The Lon protease temporally restricts polar cell differentiation events during the Caulobacter cell cycle. *Elife* **10**, doi:10.7554/eLife.73875 (2021).

7 Jonas, K., Liu, J., Chien, P. & Laub, M. T. Proteotoxic stress induces a cell-cycle arrest by stimulating Lon to degrade the replication initiator DnaA. *Cell* **154**, 623-636, doi:10.1016/j.cell.2013.06.034 (2013).

8 Holmberg, M. A., Gowda, N. K. & Andreasson, C. A versatile bacterial expression vector designed for single-step cloning of multiple DNA fragments using homologous recombination. *Protein Expr Purif* **98**, 38-45, doi:10.1016/j.pep.2014.03.002 (2014).

9 Gora, K. G. *et al.* A cell-type-specific protein-protein interaction modulates transcriptional activity of a master regulator in Caulobacter crescentus. *Molecular cell* **39**, 455-467, doi:10.1016/j.molcel.2010.06.024 (2010).

10 Wohlever, M. L., Nager, A. R., Baker, T. A. & Sauer, R. T. Engineering fluorescent protein substrates for the AAA+ Lon protease. *Protein Eng Des Sel* **26**, 299-305, doi:10.1093/protein/gzs105 (2013).
